# Deep Blood Proteomics Identifies over 12,000 Proteins Providing Valuable Information about the State of the Human Body

**DOI:** 10.1101/2025.05.15.654311

**Authors:** Zhenyu Sun, T. Mamie Lih, Trung Hoàng, Shao-Yung Chen, Joanne Xu, Ding Chiao Lin, Yuefan Wang, Jongmin Woo, Yuanyu Huang, Lijun Chen, Hongyi Liu, Marissa Alpern, Jadranka Milosevic, Hong-Zhang He, Raghothama Chaerkady, Qing Wang, Hui Zhang

## Abstract

Blood is a valuable resource for clinical research, offering insight into physiological and pathological states. However, the specific proteins detectable in blood and the optimal proteomic methods for their detection have not been rigorously investigated and documented. To address this, we conducted various blood proteomic strategies, including directly blood proteomic analysis, high-abundance protein depletion, low-abundance protein enrichment, and extracellular vesicle enrichment using data-independent acquisition or targeted proteomics. These approaches identified 11,679 protein groups in plasma from healthy individuals. In 136 pancreatic ductal adenocarcinoma whole blood samples, 6,956 protein groups were found, including 678 not seen in healthy samples, expanding the total to 12,357 blood proteins. This represents the most comprehensive blood proteome to date. To support broader access and analysis, we developed the Human Blood Proteome (HuBP) database, detailing protein detectability, abundance, and reproducibility across workflows, sample types, and disease contexts.

## Introduction

Blood is a relatively stable and easily accessible specimen in clinical applications, serving as a window into the physiological and pathological statuses ^1, 2^. It contains a rich diversity of protein, providing a distinct snapshot of the current functionality of the circulatory system and all tissues and organs in contact with blood ^3 4 5^. Typically, three types of blood samples are collected for clinical analysis: plasma, serum, and whole blood, with plasma and serum being the most used.

Mass spectrometry (MS)-based proteomics has emerged as a leading technology for exploring blood proteome due to its unparalleled specificity and unbiased ability to identify and quantify proteins ^6, 7 8^. However, MS analysis of blood proteome faces significant challenges due to the wide dynamic range of protein abundance, which spans up to 12 orders of magnitude ^9, 10^. This range complicates the detection of low-abundance proteins that are derived from disease state of tissues or organs, necessitating strategies to improve sensitivity ^3, 11^.

Approximately 1,000 protein groups can be detected by direct blood proteomic analysis using liquid chromatography-tandem mass spectrometry (LC-MS/MS) ^12^. To further increase depth, several sample preparation methods have been developed to enhance blood proteome coverage. For example, immunoaffinity-based depletion of high-abundance proteins can identify and quantify 1,557 protein groups from serum samples. Recent strategies for capturing low abundance proteins have also shown promise in enhancing blood proteome coverage ^13, 14^. Enriching extracellular vesicles (EVs) from plasma can increase the number of identified protein groups to up to 4,000 ^15^. In addition, glycosylation-specific enrichment techniques selectively capture glycoproteins, which are often low in abundance. A total of 892 glycopeptides derived from 141 glycoproteins were detected as highly expressed in pancreatic ductal adenocarcinoma (PDAC) serum samples ^16^. Nanoparticle interaction-based enrichment can also reduce the dynamic range of protein abundances, enabling the detection of up to 5,000 protein groups in plasma proteome coverage ^17^. Complete360® pipeline was developed for plasma characterization, and approximately 10,000 protein groups could be detected in plasma.

Despite significant advancements in different blood proteomic technologies, several limitations remain. While depletion columns effectively remove high-abundance proteins, they may inadvertently eliminate associated binding partners ^18^. Enrichment strategies for low abundance proteins are effective in improving blood proteome coverage, but they often face challenges with the reproducibility of blood protein identification and quantification. Additionally, the choice of sample type, plasma, serum, or whole blood, introduces further variability, as each matrix exhibits a distinct proteomic composition that affects the detection, quantification, and comparative analysis of blood-related proteins. Disease samples add another layer of complexity, with disease progression altering protein expression and enabling the identification of disease-specific proteins.

In this study, we conducted blood proteomic analysis using seven complementary strategies: direct analysis (Direct), High Select Top 14 Abundant Protein Depletion Spin Columns (Thermo), ENRICH-iST (PreOmics), the Proteograph XT workflow (Seer), the Complete360® targeted proteomics (Complete Omics), Solid-Phase Extraction of Glycopeptides (SPEG) ^13^, and extracellular vesicle (EV) enrichment ^14^. We systematically characterized diverse blood proteomic profiles from plasma, serum, or whole blood from healthy individuals. Collectively, these analyses characterized 11,679 protein groups, achieving the largest MS-based blood proteome coverage to date. Furthermore, by applying the Proteograph XT workflow to whole blood samples from 136 patients with PDAC, one of the most lethal cancer types^19^, we identified and quantified 6,956 protein groups. Among them, 6,304 proteins overlapped with PDAC tissue proteins, including 1,622 proteins demonstrating differential expression in the tumor tissues compared to normal adjacent tissues (NATs). Compared to 11,679 blood proteins identified by different methods in healthy individuals, 678 additional protein groups were uniquely identified from PDAC blood samples, resulting in 12,357 characterized blood proteins. Our findings underscored the ability of blood proteomics to capture tumor-associated proteins in blood. To provide a systematic framework for blood protein identification across multiple workflows, sample types, and disease contexts, we developed the Human Blood Proteome (HuBP) website, which presents the most extensive MS-based blood proteome to date. The HuBP not only offers a catalog of blood-derived proteins but also detailed information on detectability and reproducibility for each blood protein using different detection strategies, blood specimen types, and disease sources. Taken together, the HuBP can serve as a valuable resource for optimizing experimental designs in comprehensive blood proteomics or targeted analysis of disease-specific proteins for clinical applications.

## Results

### Protein characterization using multiple blood proteomic strategies

To detect tissue proteins in healthy and diseased states in blood, several methods have been developed to overcome the challenges of the high dynamic range of protein abundance. However, which proteins can be detected in blood and by which blood proteomic method have not been rigorously investigated. In this study, we characterized plasma proteome using seven different methods as follows: direct analysis (Direct), High Select Top 14 Abundant Protein Depletion Spin Columns (Thermo), ENRICH-iST (PreOmics), the fully automated Proteograph XT workflow (Seer), the Complete360® pipeline (Complete omics), glycosylation enrichment (SPEG), and EV enrichment method. The Seer method provides two types of nanoparticles, referred to as nanoparticle A(Seer_NPA) and nanoparticle B (Seer_NPB)^17^. Pooled human plasma samples from healthy individuals were divided into 32 aliquots, with each method performed in four replicates.

The number of protein groups identified by each method was shown in **Figure 1A**. All samples, except for the Seer method, involved manual sample preparation, while the Seer method utilized an automation sample preparation process. Direct analysis of plasma using tryptic digestion followed by DIA-MS identified 769 proteins. Depletion using the Thermo kit increased the identified protein groups to 1,434 proteins. Enrichment strategies further enhanced protein detection, with 2,328 proteins identified using the PreOmics kit, 3,573 proteins via EV enrichment, and 4,701 proteins via the Seer method (4,427 from nanoparticle A and 4,463 from nanoparticle B). SPEG identified 329 proteins. Finally, the targeted proteomic method (Complete omics) characterized a total of 9,977 proteins. Across all seven workflows, a total of 11,035 protein groups were identified from plasma, all of which were matched to the Human Protein Atlas (https://www.proteinatlas.org/humanproteome/blood), which includes 4,193 protein groups identified by MS. **(Table S1)**.

**Figure 1.**
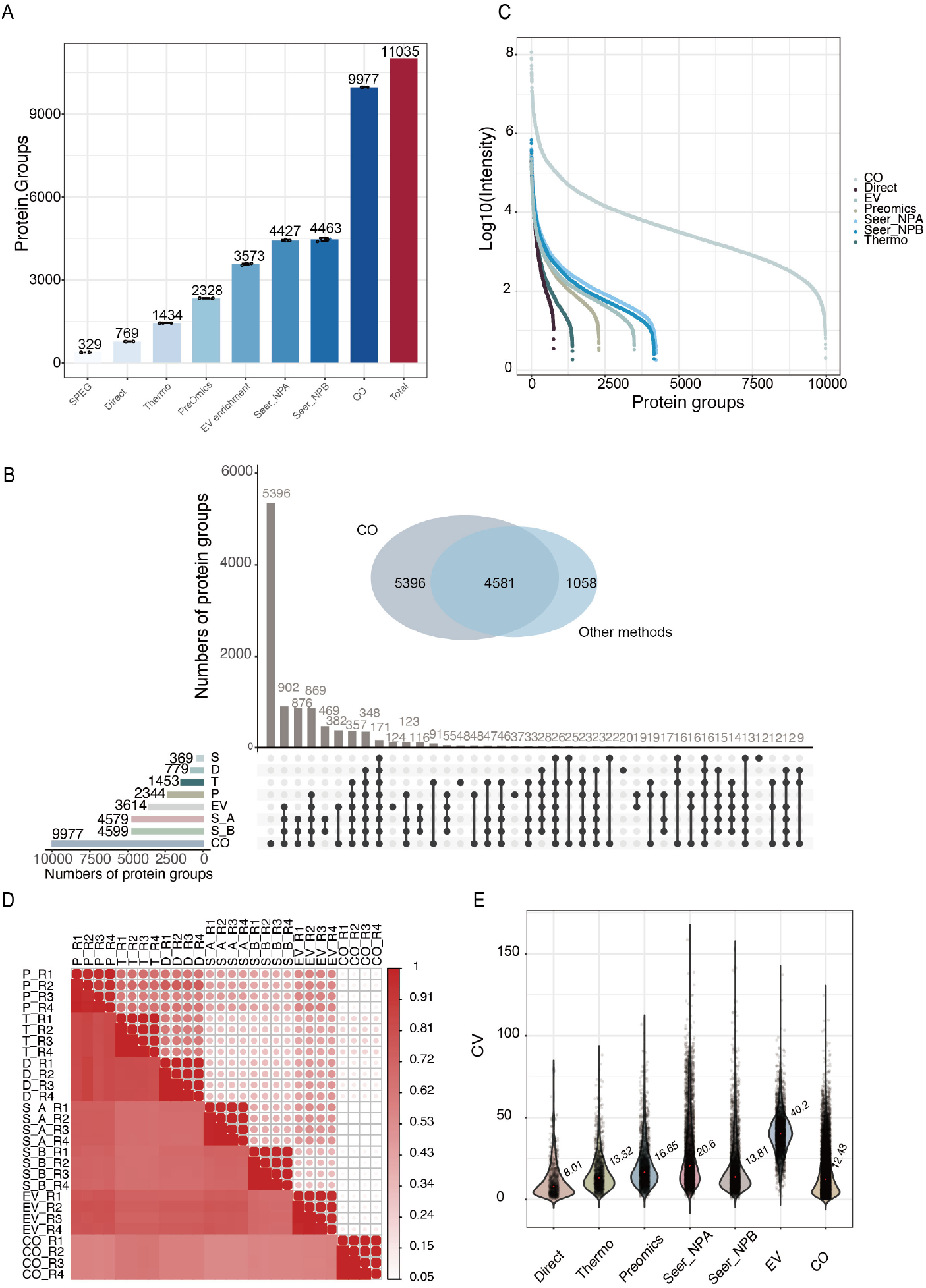
Plasma proteins identified and quantified by different blood proteomic workflows. **A** The numbers of protein groups identified across seven sample preparation methods, including SPEG, Direct, Thermo kit, PreOmics, EV enrichment, Seer method (Seer_NPA and Seer_NPB), and the Complete360®omic assay (CO). Each bar displays the mean value with individual data points, representing 4 technical replicates. “Total” represents the union of protein groups identified across all plasma workflows; **B** Unique and shared protein groups identified across the workflows, and the inset venn diagram illustrates the overlap between protein groups identified using the Complete360® assay (CO) and other six methods, including SPEG (S), Direct (D), Thermo kit (T), PreOmics (P), EV enrichment (E), Seer_NPA (S_A), and Seer_NPB (S_B). **C** Distribution of protein intensities across seven methods; **D** Pairwise correlation analysis for seven methods, including SPEG (S), Direct (D), Thermo kit (T), PreOmics (P), EV enrichment (E), Seer method (S_A and S_B), and the Complete360®omic assay (CO). “R” represents the replicate. **E** The distribution of coefficient of variation (CV) across protein groups identified by each method. Each dot represents the CV of an individual protein quantified across replicates. The mean CV for each method is labeled within the corresponding violin.

For the EV enrichment strategy, 3,614 protein groups were identified. Vesiclepedia is a widely used database of EV-associated proteins identified by various MS-based methods, and comparison with our dataset revealed an overlap of 3,126 proteins, accounting for 37% of the Vesiclepedia entries ^20^ **(Figure S1A)**. Notably, all 100 top EV marker proteins listed in Vesiclepedia were detected in our EV dataset **(Figure S1B)**. Gene Ontology (GO) analysis revealed that the primary cellular components represented were the cytoplasm, exosomes, and lysosomes **(Figure S1C)**. Glycopeptide enrichment was performed using SPEG ^13^, in which peptides were conjugated on beads, followed by the release of N-glycopeptides using PNGase F. This process identified 1,826 peptides corresponding to 329 proteins **(Figure S1D)**.

We analyzed the overlap of protein groups identified across the seven methods. A total of 171 protein groups were detected by all methods. Conversely, unique protein identifications included 20 proteins by the direct method, 48 by the Thermo method, 37 by PreOmics method, 469 by the Seer method, and 5,396 by the targeted blood proteomic analysis **(Figure 1B)**. Plasma samples remain challenging due to the broad dynamic range of protein abundance, which complicates the detection of low abundance proteins using LC–MS/MS. This variability is reflected in significant differences in protein detection across different methods.

We further characterized the abundance distribution of proteins identified by each method. The identified plasma protein abundance spanned eight orders of magnitude, with each method capturing a distinct range of mass spectrometric abundance **(Figure 1C)**. Large cohort studies require robust and reproducible workflows. Thus, we compared the quantified intensity of protein groups to assess the reproducibility both within individual replicates and across different workflows. We found that the correlation coefficients among the technical replicates for each method all exceeded 0.89, demonstrating high reproducibility within individual approaches **(Figure 1D)**. In contrast, correlations of the seven methods were relatively low, indicating variability in protein quantification across workflows. Finally, we assessed the coefficient of variation (CV) for each method across four replicate experiments. All methods, except for EV enrichment, demonstrated a median CV below 15%, further highlighting the reproducibility of the workflows **(Figure 1E)**.

### Blood proteomic analyses from different sample types including plasma, serum, and whole blood

To investigate the blood proteins from different blood sample types, we employed nanoparticle enrichment methods using Seer method to different clinical blood sample types, including serum and whole blood. Enrichment significantly increased the number of identified proteins in both serum and whole blood samples. Specifically, it resulted in a 4.8-fold increase in serum and a 3.4-fold increase in whole blood, compared to unenriched samples. The average number of identified proteins across three technical replicates was 3,115 for serum, 4,672 for plasma, and 5,647 for whole blood **(Figure 2A, Table S1)**. Notably, the enriched fraction constitutes most identified proteins, while the unenriched samples account for a smaller proportion **(Figure S2)**. Combining these results with previously identified proteins from various plasma analysis methods, a total of 11,679 detectable protein groups were cataloged from plasma, serum, and whole blood samples, representing the largest blood proteomic dataset reported to date. Protein overlap analysis among serum, plasma and whole blood samples showed that 41% of proteins were identified across all three sample types, while 5.3% were uniquely identified in plasma and 2.4% were uniquely identified in serum. Notably, 22% of proteins were uniquely identified in whole blood samples, emphasizing its potential for clinical proteomics and its advantages in capturing a broader proteome **(Figure 2B)**. To assess reproducibility, we performed three technical replicates for each sample type and analyzed their protein abundance. The intra-replicate correlation coefficients were all greater than 0.98, indicating high reproducibility for individual blood proteomic analyses of plasma, serum, and whole blood samples **(Figure 2C)**. Additionally, we calculated the CV for protein abundance in each sample type, with whole blood exhibiting the lowest CV compared to serum and plasma **(Figure 2D)**.

**Figure 2.**
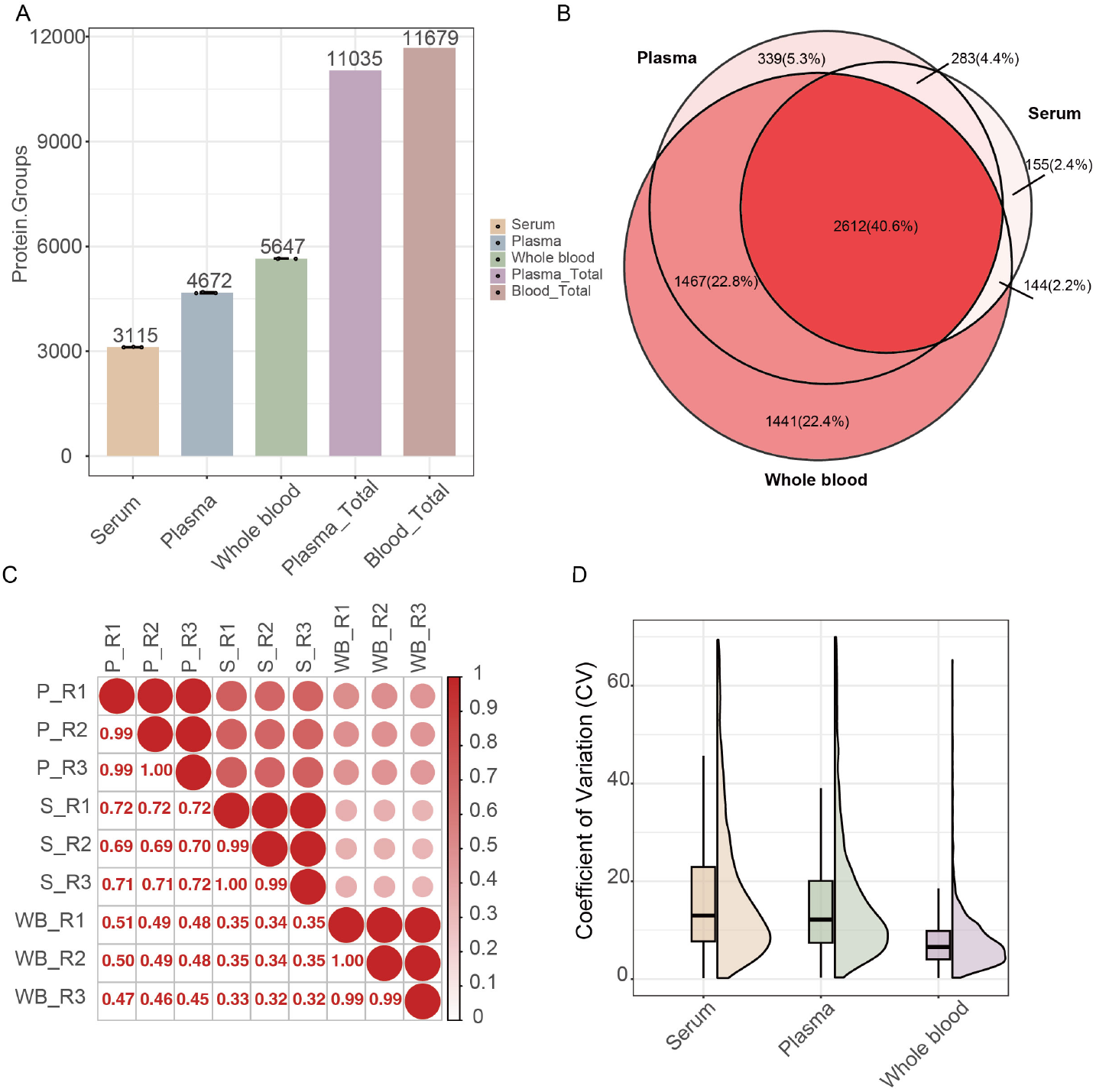
Blood protein characterization using serum, plasma, and whole blood. **A** The number of protein groups identified across serum, plasma, and whole blood using direct analysis and Seer method. Each bar displays the mean value with individual data points, representing three technical replicates. “Plasma_Total” represents the union of protein groups identified across all plasma workflows, “Blood_Total” represents the union across all blood-derived sample types including serum, plasma, and whole blood; **B** The overlap of identified proteins among serum, plasma, and whole blood samples; **C** Pairwise correlation analysis between replicates across serum (S), plasma (P), and whole blood (WB); **D** The CV distribution of each sample.

### PDAC-associated tissue proteins detected in whole blood

PDAC is often diagnosed at the advanced stage when surgical intervention is no longer feasible, making it one of the most lethal cancer types ^21^. Therefore, there is an urgent need to characterize PDAC-specific protein biomarkers for early detection and diagnosis. Early detection of PDAC through minimally invasive methods, such as blood tests, could significantly improve clinical outcomes. Utilizing the Seer method, we analyzed 136 whole blood samples from PDAC patients, identifying a total of 6,956 protein groups across all samples (**Table S1, Table S2**). On average, 6,288 protein groups were identified per sample **(Figure S3A)**. Compared to the 11,679 blood proteins identified from normal blood samples (referred to as Sample type), 678 proteins were uniquely identified from PDAC patient blood samples, resulting in 12,357 blood proteins identified from blood samples (referred to as Disease). The distribution of protein intensities, as shown in **Figure S3B**, further supports the robustness and comprehensive coverage of the dataset. In our recent study of pancreatic cancer tissues, we identified 10,183 tissue proteins from 104 PDAC tumor tissues with *KRAS* variant allele fraction (VAF) ≥ 0.075 (referred to as high-purity PDACs) and 43 normal adjacent to tumors (NATs) (**Table S1**), of which 2,441 were differentially expressed proteins (DEPs) in PDAC relative to NAT ^22^. Among these tissue proteins, 8,488 proteins (83.3%) and 8,010 proteins (78.7%) were identified from the 12,357 and 11,679 blood proteins of Disease and Sample type, respectively **(Figure 3A, Table S3)**. Among the 104 high-purity PDACs, 100 had case-matched whole blood samples. A total of 1,622 DEPs in PDACs compared to NATs from the tissues were also detected in the whole blood (**Figures 3B, Table S4**). When we included additional 35 PDAC tissues whose *KRAS* VAF < 0.075 (referred to as low-purity PDACs) that also had case-matched blood samples (**Table S1**), we detected 8,489 tissue proteins in blood (**Figure S3C, Table S1**). Similar to our previous studies, we first focused on tumors with sufficient neoplastic cellularity in the current study ^4^. The 35 low-purity PDAC tissues were initially excluded from the downstream analysis, resulted in 1,622 DEPs PDAC-associated tissue proteins.

**Figure 3.**
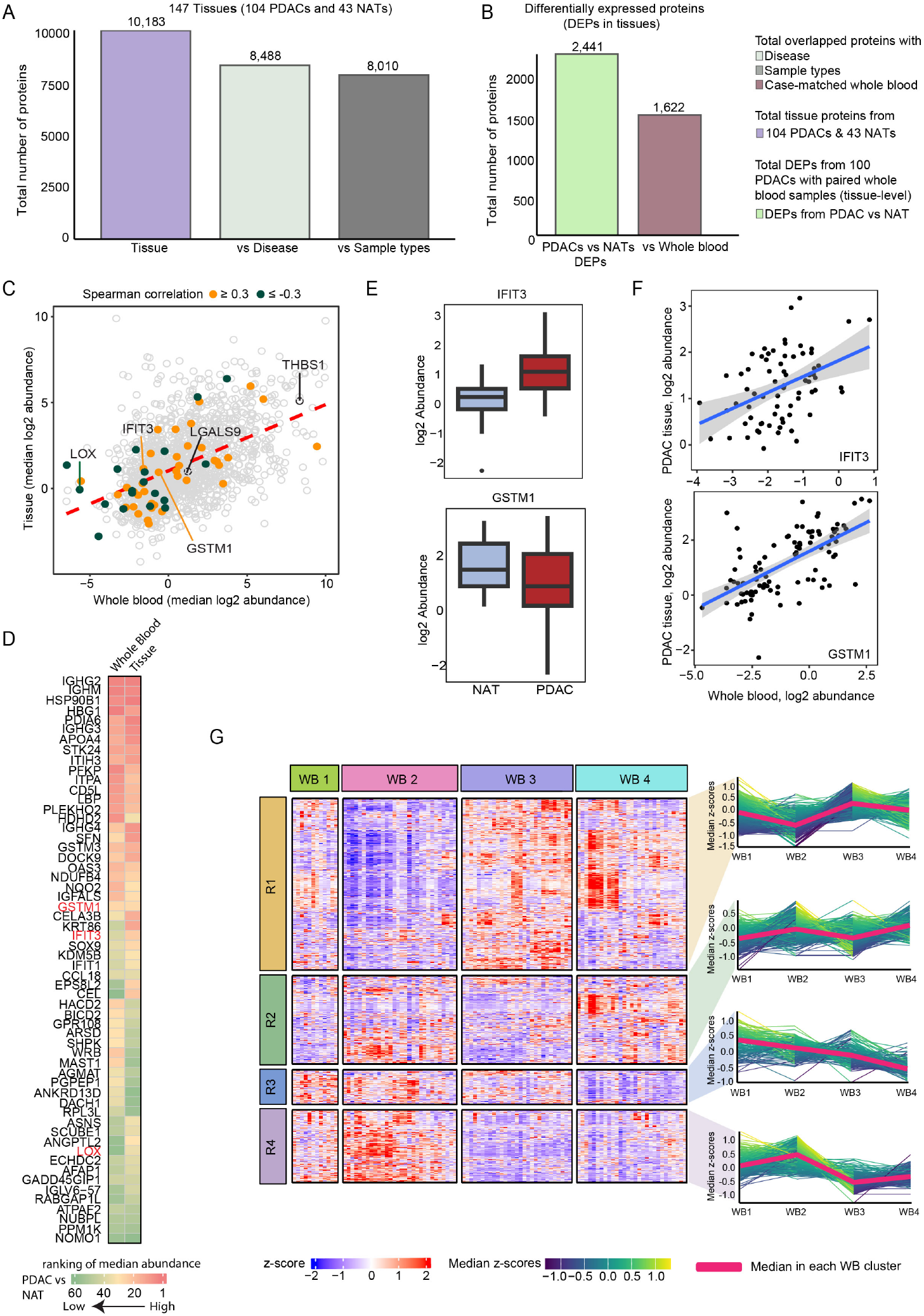
Proteomic characterization in pancreatic ductal adenocarcinoma (PDAC) tissue and case-matched whole blood samples. **A** Detection of tissue proteins in blood. A total of 11,035 blood proteins identified from pooled control sample were categorized as the “Sample types” group. Additional 136 whole blood samples from PDAC patients were analyzed, identifying a total of 12,357 proteins from both PDAC whole blood and different sample types; these were categorized as the “Disease” group. **B** Detection of differentially expressed tissue proteins (DEPs) in case-matched whole blood samples. These DEPs are considered as PDAC-associated tissue proteins. **C** Abundance association between tissue and whole blood for PDAC-associated tissue protein. Proteins with moderate Spearman correlation ≥ 0.3 (orange) or ≤ -0.3 (green) are highlighted. **D** Abundance rank distribution between PDAC tissues and case-matched whole blood samples based on the PDAC-associated tissue proteins with moderate spearman correlation. **E** Expression profiles of PDAC-associated tissue proteins, IFIT3 and GSTM1, as examples. **F** Comparison of protein abundance levels in tissue and whole blood for IFIT3 and GSTM1 as selected example proteins. **G** Hierarchical clustering of PDAC-associated tissue proteins in 100 whole blood samples with case-matched PDAC tissues. Line plots showing protein abundance profiles across sample clusters (WB1 to WB4), based on row clusters (R1 to R4).

Next, we assessed the correlation between expression profiles of PDAC-associated proteins in tissue and whole blood samples. We found that 58 proteins showed correlation of protein expression profile between tissue and whole blood with spearman correlation ≥0.3 (positively correlated) and ≤ -0.3 (negatively correlated), respectively (**Figure 3C, Table S5**). Ranking the 58 PDAC-associated proteins based on their median abundance across the 100 high-purity PDACs showed similar patterns between tumor tissues and whole blood (**Figure 3D**). Moreover, PDAC-associated tissue proteins that were identified in the whole blood could be candidates for PDAC detection using blood tests. For instance, interferon-induced protein with tetratricopeptide repeats (IFIT3) was overexpressed in PDACs relative to NATs (**Figure 3E**). A moderate positive correlation (spearman correlation of 0.4) was found between tumor tissues and whole blood (**Figure 3F**). A study has shown that upregulation of IFIT3 promotes tumor growth in pancreatic cancer ^23^. Glutathione S-transferase Mu 1 (GSTM1) was downregulated in PDACs compared to NATs, but we still observed a good correlation between its abundance in tumor tissues and whole blood samples (Spearman correlation of 0.61) (**Figures 3E-F**). On the other hand, proteins with their tumor tissue abundances negatively correlated with their abundances in whole blood may indicate that these proteins predominately expressed in tumor tissues instead of in the blood. For instance, protein-lysine 6-oxidase (LOX) was upregulated in PDACs compared to NATs, but its abundance in tumor tissues was negatively correlated with its abundance in whole blood (**Figures S3D-E**). Given that inhibiting LOX can induce tumor necrosis, it may be better characterized as a promising therapeutic target rather than a potential marker for blood testing^24^. Of note, PDAC-associated tissue proteins that had correlation outside our defined cutoffs could still play a key biological role in pancreatic cancer. For instance, thrombospondin-1 (THBS1) was detected in the 100 high-purty PDACs with higher abundance than NATs (**Figure S3F**), which was also previously identified as upregulated in tissues and serum from the patients with PDAC compared to normal tissues^25^. Galectin-9 (LGALS9) was also upregulated in PDACs (**Figure S3F**), and it could be useful for PDAC detection^26^. Furthermore, 4 distinct protein clusters (WB1 to WB4) and 4 distinct row clusters (R1 to R4) were derived based on the 1,541 PDAC-associated tissue proteins with less than 50% missing values across the 100 whole blood samples with case-matched PDAC tissues (**Figure 3G, Table S6**). These data showed that different clusters of proteins were associated with different patient subtypes, which could be used for the detection of PDACs using subtype-specific blood proteins. The heterogeneity of PDACs is also observed from previous PDAC studies by genomics, epigenomics, transcriptomics, and proteomics ^4, 16^. In addition, each row cluster was enriched with different KEGG pathways (**Figure S3G, Table S7**). For example, R2 was enriched with pathways such as, leukocyte transendothelial migration, lysosome, and carbon metabolism. On the other hand, glycolysis/gluconeogenesis and insulin signaling pathways were enriched in R4. Taken together, understanding the distribution and detectability of PDAC-associated tissue proteins in the case-matched whole blood samples could lead to potential clinical utilities of using these tissue proteins in the detection of PDAC from blood.

### Atlas of Human Blood Proteomics Development

To present the most extensive MS-based blood proteome to date as a public accessible resource for blood proteomics research, we developed the Human Blood Proteome (https://protein-notebook.streamlit.app, HuBP), a comprehensive and publicly accessible resource, encompassing data derived from diverse experimental methods, biological sample types, and disease states. This dataset represents the largest and most detailed blood proteomic resource reported to date **(Figure 4A)**. HuBP allows users to input single or multiple UniProt identifiers of proteins to generate information on which methods these proteins can be detected. It also provides plots on the corresponding MS intensities and the reproducibility performance of each method, making it a powerful tool for exploring protein detection and characterization in human blood samples. A total of 11,035 plasma proteins were identified through seven distinct experimental techniques, including CO, Seer, PreOmics, Thermo, Direct, EV enrichment, and SPEG methods. Each method provided uniquely contributed to the overall depth and coverage of the proteome.

**Figure 4.**
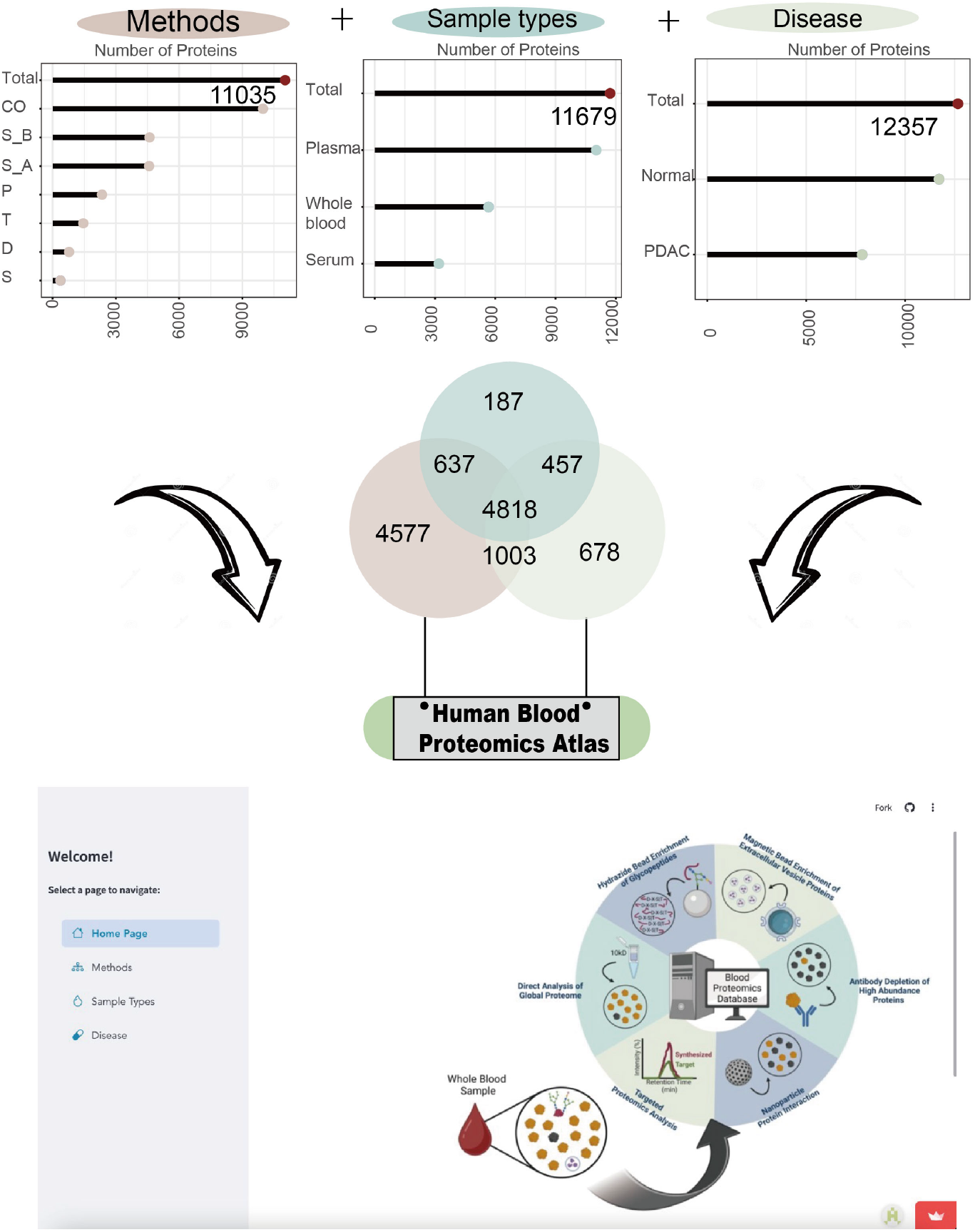
Comprehensive Overview of the. Human Blood Proteome. A total of 11,035 proteins were identified using various approaches, including SPEG (S), Direct (D), Thermo kit (T), PreOmics (P), EV enrichment (E), Seer method (S_A and S_B), and the Complete360® assay (CO); A total of 11,679 proteins were identified from plasma, whole blood, and serum samples; A total of 12,357 proteins were identified across normal and disease conditions, including PDAC.

Through analyzing plasma, serum, and whole blood samples from healthy individuals, a combined total of 11,679 proteins were identified, with each matrix contributing a unique subset of proteins. This collective data provides a representative profile of the normal blood proteome. Furthermore, integrating data from disease conditions, including PDAC, expanded the dataset to 12,357 proteins, offering insights into both shared and disease-specific protein signatures.

The HuBP provides a comprehensive resource for researchers, offering detailed insights into each protein’s detectability across analytical methods, association with specific clinical sample types, and relevance to disease states. By including data on signal intensity and quantification reproducibility, the HuBP allows users to make informed decisions when selecting analytical approaches, identifying the optimal clinical sample type for detection, or investigating links between proteins and specific diseases. For instance, the immunoglobulin lambda variable 5-37 (IGLV5-37, A0A075B6J1) is part of the structure of antibodies, which are produced by B cells and play a critical role in the immune response by binding to antigens. This protein was only identified by Direct, PreOmics and CO methods and was not captured by the other methods **(Figure S4)**. Notably, it exhibited high MS intensity and a CV below 10% with PreOmics method, indicating both strong detectability and precise quantification. This integrated framework makes the HuBP an invaluable tool for advancing basic research and translational applications in proteomics. Detailed user guidance is included in the supporting information, facilitating accessibility and effective utilization of this atlas.

## Discussion

In this study, we developed Human Blood Proteome (HuBP), a comprehensive blood proteomics resource consisting of 12,357 proteins identified and quantified across multiple experimental techniques, clinical sample types, and pancreatic ductal adenocarcinoma (PDAC) disease states. To the best of our knowledge, this represents the most extensive MS-based blood proteomic dataset reported to date.

A systematic evaluation of various sample preparation strategies revealed distinct advantages and applications in enhancing the depth, specificity, and reproducibility of plasma proteomics. The direct analysis strategy, while achieving the lowest protein identification numbers, requires minimal plasma (could less than 1µL) and employs a simple workflow, making it particularly suitable for studies involving limited sample volumes. The Thermo method efficiently removes the top 14 high-abundance proteins within 30 minutes and achieved lower CV compared to nanoparticle-based methods. In contrast, the Complete360®omic assay yielded the highest number of plasma protein groups but exhibited higher costs and lower scalability. The nanoparticle-based enrichment method demonstrated high sensitivity, particularly in detecting low-abundance proteins that are typically missed by conventional workflows. These capabilities are especially valuable in clinical proteomics, where comprehensive detection of low-abundance proteins is essential for biomarker discovery and disease characterization. Moreover, the automated Proteograph XT workflow from Seer improved reproducibility compared to manual processing, reinforcing the role of automation in enhancing data quality.

Specialized enrichment strategies further broadened proteome coverage. The EV enrichment method from Captis enabled the identification of a significant number of vesicle-associated proteins, including 3,126 proteins overlapping with the Vesiclepedia database. This supports its utility for EV proteomics, which is critical for studying intercellular communication, disease mechanisms, and biomarker discovery ^27^. Similarly, the SPEG enrichment method effectively captured glycopeptides containing the N-X-S/T motif (where X ≠ P) to facilitate in-depth glycoproteomic profiling. Given the essential roles that glycoproteins play in physiological regulation and disease progression, specialized enrichment strategies like SPEG may be particularly suited for characterizing glycosylation changes in blood ^28^.

Beyond method-specific evaluations, we also compared different blood sample types, plasma, serum, and whole blood, to assess their contribution to proteome coverage. Each sample type exhibited distinct proteomic profiles. Notably, 22% of proteins were unique to whole blood, likely reflecting the presence of cellular components absent in plasma or serum. These findings underscore the added value of whole blood in clinical proteomics, particularly for studies investigating cellular processes and disease mechanisms.

To further explore the clinical relevance of our dataset, we analyzed proteomic profiles from 136 PDAC whole blood samples. Compared to proteins identified in healthy individuals, 678 proteins were uniquely associated with PDAC, representing a valuable pool of candidate biomarkers. Notably, 100 of these PDAC whole blood samples had case-matched tumor tissues with sufficient neoplastic purity that were previously analyzed in our recent study. Comparative analysis revealed that many PDAC-associated tissue proteins exhibited consistent expression trends in whole blood, suggesting that blood-based proteomics can effectively capture tumor-derived molecular signals. These findings support the potential of whole blood as a minimally invasive platform for PDAC detection and disease monitoring.

Collectively, the HuBP offers a valuable resource for both basic research and translational applications in proteomics. By providing detailed data on protein detectability, signal intensity, and quantification reproducibility across diverse workflows, clinical sample types and disease states, the HuBP enables researchers to make informed decisions about analytical approaches and sample selection.

## Conclusion

This study presents a significant advancement in blood proteomics by addressing the critical question of which proteins can be detected in blood and through which proteomic methods, an area that has not been rigorously investigated or systematically documented. Collectively, 11,679 protein groups were characterized across seven different methods in serum, plasma, and whole blood samples, representing the largest MS-based blood proteome coverage to date. Furthermore, the analysis of 136 whole blood samples from PDAC patients highlighted the utility of whole blood as a minimally invasive alternative to tissue samples for biomarker discovery. A total of 6,956 protein groups were identified in PDAC samples, including 678 proteins uniquely detected in cancer cases. By examining the 100 whole blood samples with case-matched high-purity PDAC tumor tissues, we identified 58 PDAC-associated tissue proteins whose abundances in tumors correlated better with their abundances in the whole blood. These findings demonstrate the connection between the whole blood and tumor proteomes, emphasizing the potential of utilizing whole blood in clinical proteomics and disease biomarker discovery. The development of the HuBP, comprising of 12,357 blood proteins across multiple experimental techniques, sample types, and PDAC disease states, provides an invaluable resource for the research community. By enabling comprehensive proteome coverage and supporting biomarker discovery, this work contributes to a deeper understanding of human health and disease, paving the way for the development of novel diagnostic tools and therapeutic strategies.

## Methods

### Subject details

The plasma, serum, and whole blood samples were purchased from Innovative Research, Inc. The 136 PDAC whole blood samples were from the Clinical Proteomic Tumor Analysis Consortium (CPTAC). Tumor tissues and NATs from PDAC patients were from our previous study ^29^. Institutional review boards at tissue source sites, reviewed protocols and consent documentation adhering to the CPTAC guidelines for study participation. Both PDAC whole blood samples and tissue samples used in this study were prospectively collected for the CPTAC project. Of note, we only used the expression matrix from the previous study for the tissue samples ^22^.

### Non-treatment plasma sample preparation

Take 10 μL of serum and dilute it with 190 μL of 8 M urea before transferring it into a 10 kDa ultrafiltration tube. Centrifuge at 14,000 rpm for 50 minutes at 8°C to initiate ultrafiltration and urea exchange. Then, add 200 μL of 50 mM Tris-HCl buffer, centrifuging at 14,000 rpm for 50 minutes at 4°C, and repeat this step three times to ensure thorough buffer exchange. Invert the ultrafiltration tube into a 1.5 mL centrifuge tube and centrifuge at 1,000 rpm for 10 minutes at 4°C to collect the filtrate. Add 50 μL of 50 mM Tris-HCl solution to the ultrafiltration tube, centrifuge again at 1,000 rpm for 10 minutes at 4°C, and repeat this step twice, combining all collected filtrates to obtain the serum protein solution. Peptide concentrations were determined using NanoDrop, and 400 ng of each sample was utilized for LC-MS/MS analysis.

### Plasma sample preparation using the ENRICH-iST workflow

Plasma samples were prepared using the ENRICH-iST Kit (PreOmics, Germany) according to the manufacturer’s instructions. Briefly, 20 µL of plasma was incubated with pre-washed EN-BEADS in EN-BIND buffer at 30 °C and 1200 rpm for 30 minutes. The proteins bound to the EN-BEADS were washed three times using wash buffer. Subsequently, 50 μL of LYSE-BCT buffer was added to each tube, and samples were heated at 95 °C for 10 minutes. And then, the trypsin digestion buffer was added, and the samples were incubated at 37 °C for 3 hours with shaking at 1200 rpm. The digestion was stopped by adding the provided stop buffer, and the reaction supernatant was cleaned up using the kit’s filter cartridge. Peptides were eluted twice with 100 μL of elution buffer and combined. The peptides were then dried in a speed-vac and resuspended in 15 μL of 3% ACN(v/v)/0.1% FA (v/v). Peptide concentrations were determined using NanoDrop, and 400 ng of each sample was utilized for LC-MS/MS analysis.

### Plasma sample preparation using the Proteograph XT workflow

A total of 240 µL plasma sample was used for Proteograph XT workflow (Seer Inc., USA). The corona formation, wash, protein lysis and alkylation, digestion, and peptide cleanup were done on Proteograph XT workflow on SP100 Automation Instrument (Seer) as described [17]. The eluted peptides were dried in a speed-vac and resuspended in 3% ACN(v/v) /0.1% FA(v/v). A total of 400 ng of each sample was utilized for LC-MS/MS analysis. Serum and whole blood samples will be treated in a similar way.

### Plasma sample preparation using Thermo High-Select Top14 abundant protein depletion resin

Plasma samples were prepared using the Thermo High-Select Top14 abundant protein depletion resin according to the manufacturer’s instructions (Thermo Fisher Scientific, USA). Briefly, 10 µL of plasma was directly added to the resin slurry in the column. Incubate the mixture in the column with gentle end-over-end mixing for 10 minutes at room temperature. After incubation, snap off the bottom closure and loosen the top cap. Place the column into a collection tube and centrifuge at 1,000 × g for 2 minutes. Discard the column containing the resin. Protein will be stored at -20°C for later use.

### Isolation of extracellular vesicles isolation from human plasma

After thawing the de-identified pooled human plasma samples at 37°C, extracellular vesicles (EVs) were isolated from all samples using the modified lipid nanoprobe protocol (Lipid Nanoprobe Plasma EV Isolation Kit for Proteomic Analysis, Catalogue No. E0001P, Captis Diagnostics Inc).^30^ The EVs captured by the lipid nanoprobe were washed six times with 750 µL of 1X Phosphate Buffered Saline (PBS). Subsequently, the captured EVs were dried in a SpeedVac vacuum concentrator at 45°C for 2 hours. Finally, the dried EVs were lysed using 5% SDS.

### Extracellular vesicles protein sample preparation

S-Trap™ was used for protein digestion and followed the manufacturer’s instructions with slight modification (ProtiFi, USA) ^31^. In brief, the EV protein pellet was dissolved in 100 μL of 5% SDS/10 mM triethylammonium bicarbonate (TEAB), reduced with 120 mM TCEP, and held for 15 min at 55 °C in a thermomixer, and alkylated with 500 mM MMTS at room temperature for 10 min. The lysate was acidified with phosphoric acid. A total of 540 μL of S-Trap membrane binding and washing buffer (SBW) (90% methanol/100 mM, TEAB) was added to the solution and mixed well via pipetting. All samples including anything insoluble were transferred to the S-Trap filter via centrifugation at 10,000g for 1min. The filter was washed three times with 150 μL of SBW at 10,000g for 1 min. Trypsin and LysC mixture (∼1:10 enzyme/protein) in 30 μL of 50 mM TEAB were added to the surface of the filter and incubated for 2 h at 47 °C. The tryptic peptides were eluted via centrifugation for 1 min at 10,000g sequentially with 40 μL of TEAB, 40 μL of 0.2%TFA(v/v), and 40 μL of 50% ACN (v/v) 0.2% TFA (v/v). The eluted peptides were vacuum-dried. Peptide concentrations were determined using NanoDrop, and 400 ng of each sample was utilized for LC-MS/MS analysis.

### N-Glycopeptide sample preparation

The N-glycopeptides were enriched using the SPEG method ^13, 32^. In brief, the tryptic peptides were oxidized with 10 mM sodium periodate, with the reaction carried out at room temperature in the dark. The peptides were desalted using C18 columns. Hydrazide resin was prepared by washing it three times with 1 mL water. The reaction was incubated at room temperature overnight with gentle shaking. After that, the beads were washed three times with 1 mL of 50% ACN (v/v), 1.5 M NaCl, water, and 25 mM NH_4_HCO_3_. The resin was then resuspended in 200 μL 25 mM NH_4_HCO_3_. PNGase F was added to the solution to release N-glycopeptides from the beads. The supernatant was collected. And the supernatant was cleaned up by C18 columns for further LC-MS/MS analysis.

### SepPak C18 desalting of peptides

Peptides were desalted with C18 cartridges (Waters Corporation, USA). C18 Cartridges were conditioned in 1 mL ACN twice and balanced twice each with 1 mL 50% ACN (v/v)/0.1% FA (v/v) and 0.1% TFA (v/v). The peptides were loaded and washed in 1 mL 0.1% TFA (v/v) twice, followed by 1 mL 1% FA (v/v). and then elution using 500 μL 50% ACN (v/v)/0.1% FA (v/v) twice. The samples were dried in a vacuum centrifuge and stored at -80 °C.

### The Complete360® pipeline sample preparation

Plasma samples were analyzed using the Complete360® pipeline (Complete Omics Inc., USA). Briefly, human plasma samples were thawed and depleted of high- and medium-abundance proteins using an in-house-prepared sample preparation kit, trademarked as the Booming® kit. The remaining proteins were enzymatically digested to generate plasma peptidome samples. To enhance the detection and reproducibility of low-abundance targets, the MaxRec procedure was applied, boosting signal intensity while maintaining consistent quantification. Fractionation was performed using a high-performance liquid chromatography (HPLC) system configured with a customized loading program. This program enabled low-pH loading and high-pH fractionation, ensuring high reproducibility across samples with following LC gradient for reversed-phase C18 column: 5% to 28% acetonitrile over 75 minutes, followed by an increase to 42% over 8 minutes, and final ramp to 98% over 3 minutes; flow rate maintained at 1 mL/min. The resulting fractions were collected in a sample plate pre-quenched with selected MaxRec sequences tailored for each fraction. These fractions were sequentially analyzed using a rapid-fire mass spectrometry schedule, designed to ensure uninterrupted data collection by employing multiple in-house-prepared HPLC columns, eliminating delays associated with column reconditioning. Peptide abundances from each fraction were quantified and normalized using pre-selected internal standards. The resulting data provided normalized protein abundance values, enabling reliable comparisons of protein abundance across samples analyzed using the Complete360® pipeline.

### Liquid chromatography with tandem mass spectrometry (LC–MS/MS) analysis

An Evosep One LC system (Evosep Biosystems, Germany) was coupled to a timsTOF HT mass spectrometer (Bruker, Germany). Peptides were loaded on Evotip (Evosep Biosystems, Germany) and separated on PepSep C18 column of 15 cm x 150 µm, 1.5 µm (Bruker, Germany) in Bruker column toaster (50 °C) at a 30 SPD gradient. The data was acquired using the diaPASEF mode on timsTOF-HT with settings as follows: MS1 scan range of 100-1700 m/z; MS2 scan range of 338-1338 m/z, mass width 25.0 Da without mass overlap, 1 mobility window, 1/K0 range of 0.70-1.45 V.s/cm2, ramp time 85.0 ms. For samples prepared using the Complete360® pipeline, targeted quantitative proteomics was performed using an Agilent 6495D QqQ Mass Spectrometer (Agilent Technologies, Inc., USA) equipped with dynamic Selected Reaction Monitoring (dSRM) methods. Chromatographic separation was achieved using an in-house packed C18 column (1.7 µm, 2.1 × 30 mm) with the following LC gradient: 12% to 42% solvent B over 5.4 minutes, followed by an increase to 98% over 1 minute; flow rate maintained at 150 µL/min.

## Supporting information

Supporting information for this maunscript

## Data analysis

Spectral library was generated in Spectronaut® 19.1 (Biognosys AG, Switzerland). The Pulsar search settings for the protein search archive generation were selected full-specific tryptic digestion with up to two missed cleavages within the length range of 7– 52 amino acids. The fixed modification was on carbamidomethylation of cysteine (C). The variable modifications were on oxidation of methionine (M) and acetylation of protein N-terminal. The library generation settings followed BGS factory settings with all PSM FDR, peptide FDR and protein FDR below 0.01. For the glycosite containing peptide library generation, a variable modification from N to D was added and other settings were the same as global proteomic library generation. The proteins were identified and quantified by modified sequence, without cross-run normalization. The expression matrices of PDAC whole blood samples and tissue samples were median normalized. The abundance association was computed based on the Spearman correlation between 100 high-purity PDAC tissues and their case-matched whole blood samples.

For the targeted proteomic data, DIA-NN and FragPipe were used for processing both DDA-PASEF and DIA-PASEF datasets, ensuring comprehensive peptide and protein identification ^33 34^. CompletePeaking® Algorithm (Proprietary AI-Based Software) was utilized to train on over thousands of curated MS datasets for advanced peak picking and quantification. Employs machine learning techniques, including XGBoost, for accurate peak identification and effective noise reduction. The spectral library used were generated using full tryptic digestion, allowing up to two missed cleavages, including the following modifications: Carbamidomethylation (C) as a fixed modification; Oxidation (M) and N-terminal acetylation as variable modifications. The FDR was set as <1% at the PSM, peptide, and protein levels. Protein quantification was performed using Multi-point Normalized Protein eXpression (MNPX) values. Data normalization employed internal, endogenous stable standard biomarkers to minimize technical variation. Analytical reproducibility was maintained with a CV below 10% for high-confidence targets, ensuring clinical-grade precision.

## Data availability

The HuBP web application is accessible at https://protein-notebook.streamlit.app/. The source code, enabling further customization and reproducibility, is available at https://github.com/alvinhoang01/Protein_notebook. The mass spectrometry proteomics data have been deposited to the ProteomeXchange Consortium via the PRIDE partner repository with the dataset identifier PXD063544 ^35^. The reviewer can access the dataset by logging in to the PRIDE website using the following account details: Username: Password: kUHlrmnEhllB

## Acknowledgement

This work was supported by National Institutes of Health, National Cancer Institute (CA093900), the Clinical Proteomic Tumor Analysis Consortium (CPTAC, U24CA271079), the Early Detection Research Network (EDRN, U2CCA271895), Pancreatic Cancer Detection Consortium (PCDC, U01CA274514), and US Department of Defense, the Patrick C. Walsh Prostate Cancer Research Fund (CDMRP/PCRP, W81XWH-20–10353 and W81XWH-22-1-0680).

## Inclusion & Ethics

This study used de-identified samples from the Clinical Proteomic Tumor Analysis Consortium (CPTAC) and commercial sources under prior ethical approval. All authors met authorship criteria with roles defined before the study.

