## Supporting information for this maunscript for "Deep Blood Proteomics Identifies over 12,000 Proteins Providing Valuable Information about the State of the Human Body"

### Table of Contents

Figures ............ 3

Figure S1. Plasma proteins identified by different blood proteomic methods. ............ 3

Figure S2. The overlap identified proteins among serum, plasma, and whole blood. ... 4

Figure S3. Detection of PDAC tissue proteins in case-matched whole blood samples. 5

Figure S4. Bar plot of mean MS intensity and coefficient of variation (CV) for immunoglobulin lambda variable 5-37 (IGLV5-37, A0A075B6J1) as detected by different blood proteomic methods. ............ 7

User guide for blood protein database ............ 8

### Figures

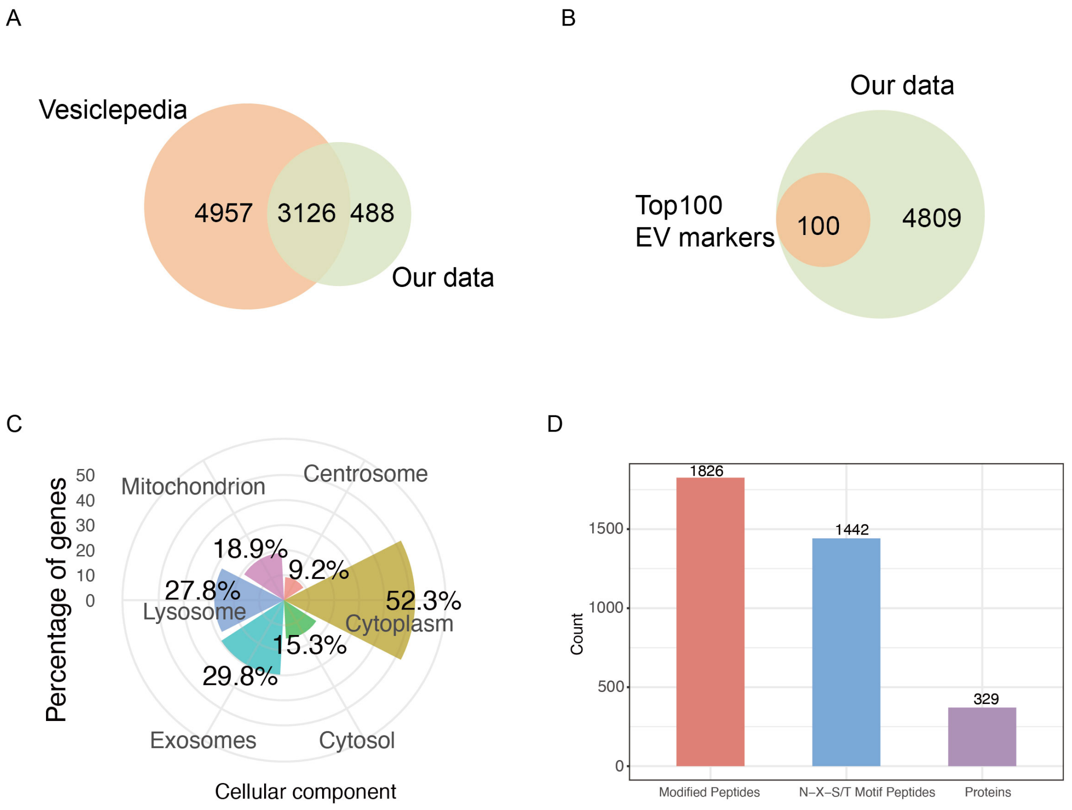


**Figure S1. Plasma proteins identified by different blood proteomic methods. A** Overlap of proteins identified from our dataset using EV method with the Vesiclepedia database. **B** The top 100 recognized EV marker proteins were all identified in our dataset using EV method. **C** GO (Gene Ontology) analysis of identified proteins based on cellular components. **D** Proteins intensity distribution curve of plasma protein abundance. Proteins only detected by the Seer method and with low intensity are highlighted in red.

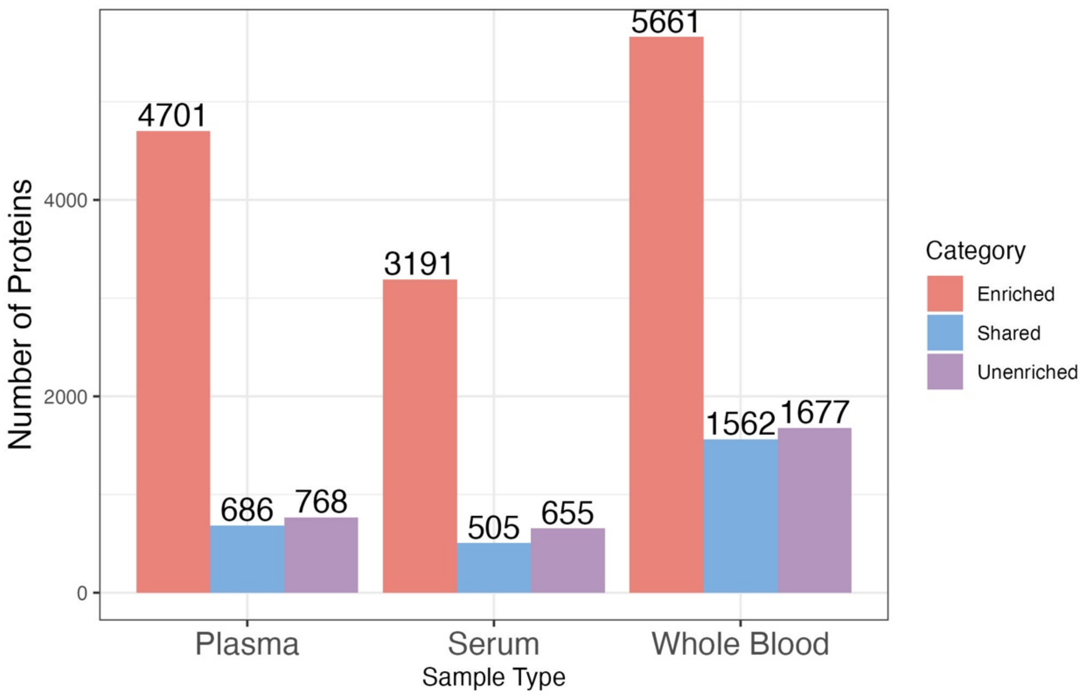


**Figure S2. The overlap identified proteins among serum, plasma, and whole blood.**

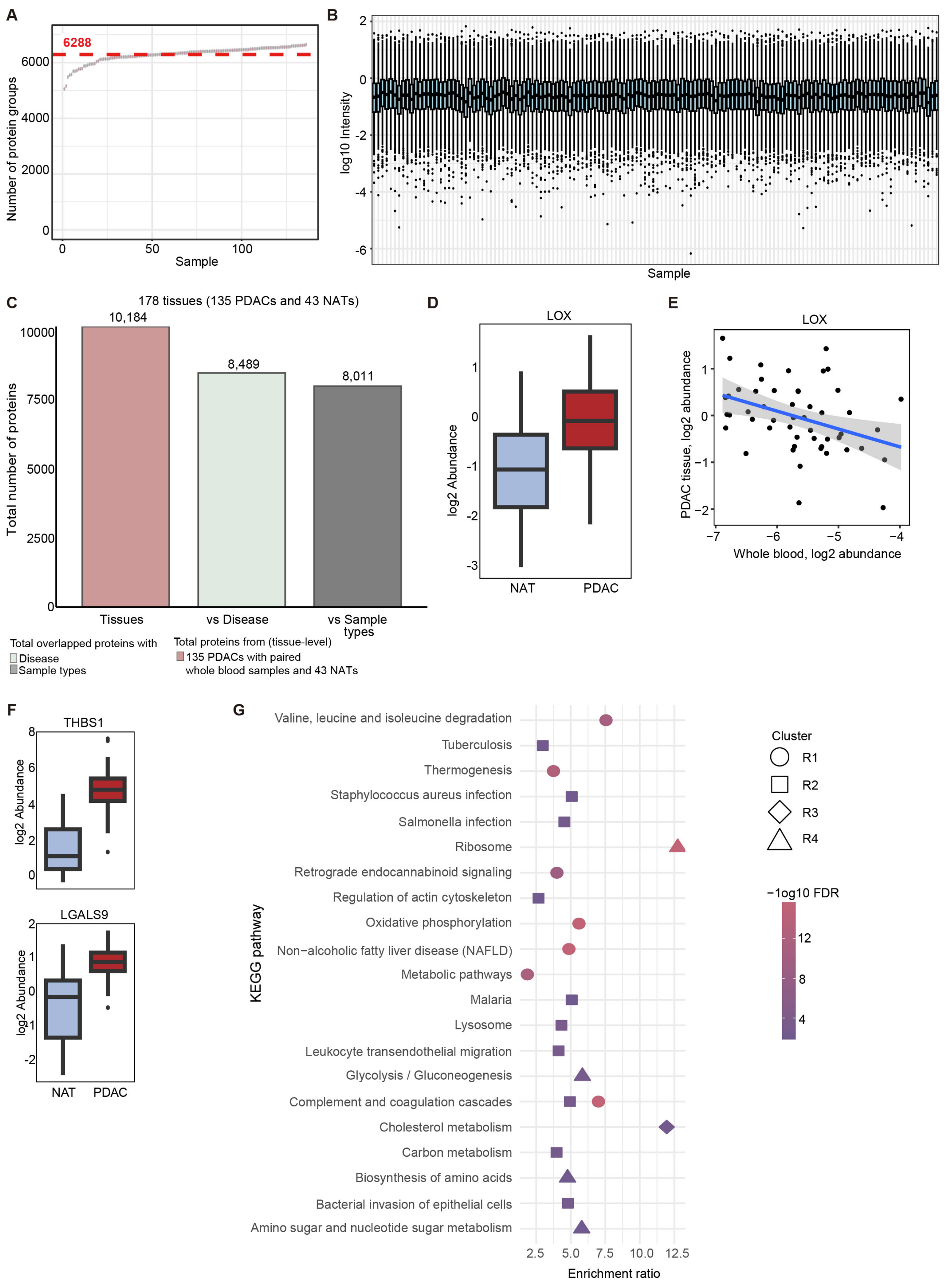


**Figure S3. Detection of PDAC tissue proteins in case-matched whole blood samples.** A The number of protein groups identified from each PDAC whole blood sample. B The distribution of total MS intensity for each patient. C Detection of tissue proteins in blood. Alongside 100 PDAC tissues with KRAS VAF ≥ 0.075 (high neoplastic cellularity), 35 additional PDAC tissues with KRAS VAF < 0.075 that had case-matched whole blood samples are included here. A total of 11,027 blood proteins identified from pooled control sample were categorized as the "Sample types" group. Additional 136 whole blood samples from PDAC patients were analyzed, identifying a total of 12,357 proteins from both PDAC whole blood and different sample types; these were categorized as the "Disease" group. **D** Expression profile of a PDAC-associated tissue protein whose abundance in PDAC tissues correlates with its abundance in whole blood. **E** Comparison of protein abundance levels in tissue and whole blood for the selected example protein. **F** Examples for demonstrating expression profiles of PDAC-associated tissue proteins (from PDACs vs NATs) whose abudnace in PDAC tissues did not show strong correlation with their abundances in whole blood according to the correlation cutoff of ≥0.3 or ≤-0.3. **G** Enriched KEGG pathways for the row clusters, R1 to R4.

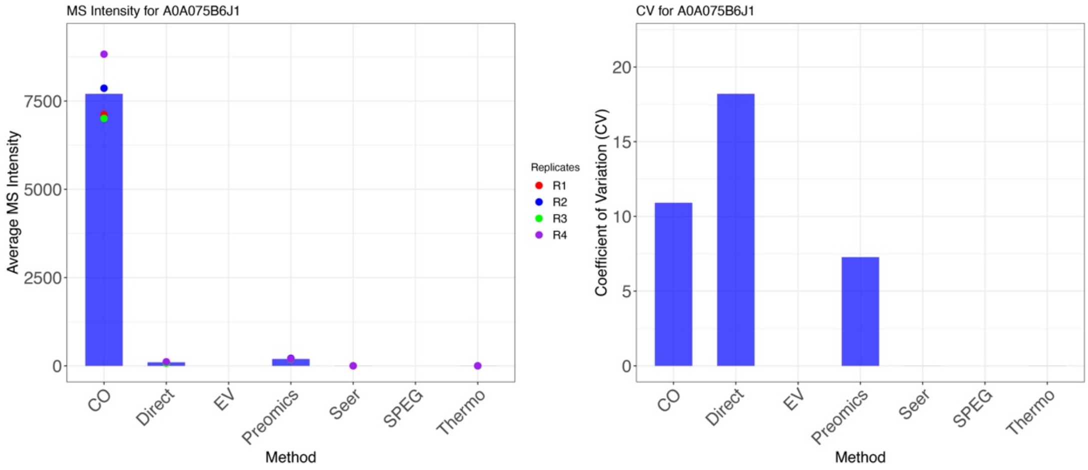


**Figure S4. Bar plot of mean MS intensity and coefficient of variation (CV) for immunoglobulin lambda variable 5-37 (IGLV5-37, A0A075B6J1) as detected by different blood proteomic methods.**

### User guide for blood protein database

This website describes the blood protein database consisting of a homepage and three main sections designed to help users explore and analyze blood proteins. Below is a detailed overview of each section:

1. **Methods**:
   This section describes the identification of protein groups in plasma using seven different approaches:
   - Non-treatment strategy (*Direct*),
   - Thermo Scientific High Select Top 14 Abundant Protein Depletion Spin Columns (*Thermo kit*),
   - Commercially available sample preparation kit ENRICH-iST (*PreOmics*),
   - Fully automated Proteograph XT workflow (*Seer*),
   - Complete360® pipeline (*CO*),
   - Glycosylation enrichment (*SPEG*), and
   - Extracellular vesicle enrichment method (*EV*).

Users can input or select UniProt IDs using the search tool. After clicking **"Search"**, the platform generates:

   - A *MS Intensity Plot*: Displays the mass spectrometry intensity for each method.
   - A *CV Plot*: Shows the coefficient of variation (CV) for each method, allowing users to evaluate both the intensity and reproducibility of the results.

2. **Sample Types**:
   This section provides information on the identification of protein groups in different biological matrices, including serum, plasma, and whole blood.
   Users can input or select UniProt IDs, similar to the *Methods* section. After clicking **"Search"**, the platform generates a presence summary across the sample types. The resulting plot indicates where the selected proteins are identified, with count numbers representing the number of identified proteins.
3. **Disease types**:
   This section focuses on the identification of protein groups in healthy controls and PDAC (pancreatic ductal adenocarcinoma). Proteins identified using different methods and sample types are used as the baseline for comparison.
   As in the previous sections, users can input or select UniProt IDs. After clicking **"Search"**, the platform generates a presence summary. The plot highlights where the selected proteins are identified in the samples, with count numbers representing the number of identified proteins.
